# Automated cell boundary and 3D nuclear segmentation of cells in suspension

**DOI:** 10.1101/632711

**Authors:** Benjamin Kesler, Guoliang Li, Alexander Thiemicke, Rohit Venkat, Gregor Neuert

## Abstract

To characterize cell types, cellular functions and intracellular processes, an understanding of the differences between individual cells is required. Although microscopy approaches have made tremendous progress in imaging cells in different contexts, the analysis of these imaging data sets is a long-standing, unsolved problem. The few robust cell segmentation approaches that exist often rely on multiple cellular markers and complex time-consuming image analysis. Recently developed deep learning approaches can address some of these challenges, but they require tremendous amounts of data and well-curated reference data sets for algorithm training. We propose an alternative experimental and computational approach, called CellDissect, in which we first optimize specimen preparation and data acquisition prior to image processing to generate high quality images that are easier to analyze computationally. By focusing on fixed suspension and dissociated adherent cells, CellDissect relies only on widefield images to identify cell boundaries and nuclear staining to automatically segment cells in two dimensions and nuclei in three dimensions. This segmentation can be performed on a desktop computer or a computing cluster for higher throughput. We compare and evaluate the accuracy of different nuclear segmentation approaches against manual expert cell segmentation for different cell lines acquired with different imaging modalities.

## Introduction

Individual cells respond to their environment, make cell fate decisions, or cause diseases when mutated. Understanding biological processes in detail at the molecular and cellular level in healthy and diseased tissue ultimately requires that cells be analyzed at single cell resolution. Over the last few decades, microscopy techniques to image single cells have improved significantly, resulting in a wealth of imaging data sets^1–4^. However, analyzing these data sets quantitatively in a high-throughput manner is still an immensely difficult and often unsolved task^5^. The lack of quantitative analysis algorithms is the ultimate bottleneck in extracting more information from microscopy images, thus hindering mechanistic understanding of single cell behavior and its relevance to physiology and disease^2,6^.

To address these limitations, efforts have been undertaken to develop image processing platforms to analyze microscopy images and segment individual cells^2,4,7^. These approaches require high-contrast fluorescent cellular markers to distinguish between the inside of the cell and the cell boundary. In many cases, cells are grown under low density adherent cell culture conditions, resulting in few cells that have very low contrast when imaged in widefield^8,9^. Furthermore, because of low-contrast images, cell segmentation in mammalian cells requires fluorescent markers to stain the cell boundary homogenously^10^. However, variability in protein expression levels, cell morphology or cell preparation for microscopy can cause heterogenous staining for an individual marker, making single-color stains insufficient for achieving homogenous staining and requiring human-assisted cell segmentation^11^.

Cell staining approaches utilizing multiple cell segmentation markers labeled with different fluorophores have been used to address this problem on the experimental side^12,13^. While the effects of heterogeneous staining can be minimized by this approach, the use of additional cell segmentation markers in different fluorescent channels reduces the number of channels available to address biological questions^12,13^. Additionally, this methodology is time-consuming due to its requirement of significant optimization and long periods of imaging. Regardless of the staining strategy, most adherent cells are grown at low density to enable cell segmentation, resulting in few cells per field of view and limited single-cell statistics. Cells in suspension are easier to segment, but concentrating cells to ensure high density during imaging to increase cell statistics is challenging.

Computationally, attempts have been made to address image analysis challenges using machine learning and deep learning approaches^3,9,14^. Such approaches rely on large and well-curated data sets to train a complex model describing features in images that identify single cells. Studies report impressive results using these approaches in cell segmentation, but it is currently an open question how transferable models generated by these approaches are to new and much smaller data sets^14^. Implementing machine learning and deep learning approaches to generate new models requires substantial expertise, making such methods extremely difficult for less-experienced users to adopt^15^.

We propose an alternative approach, called CellDissect, that overcomes these limitations both experimentally and computationally to improve image segmentation. The approach involves first optimizing sample preparation. By utilizing single-cell dissociation approaches alongside repurposed commercial “well-in-a-well” technology, CellDissect’s experimental workflow results in high-contrast widefield images with high cell density (Figure 1A). This is paired with nuclear staining, which allows for segmentation of the nucleus and assists cell boundary segmentation without the use of multiple markers or fluorescent channels. CellDissect’s nuclear and cellular segmentation algorithms then process these images with minimal user input through MATLAB or a graphical user interface (GUI) to generate highly accurate nuclear segmentation in 3D and cellular segmentation in 2D without the need for computational expertise or large, curated datasets.

**Figure 1.**
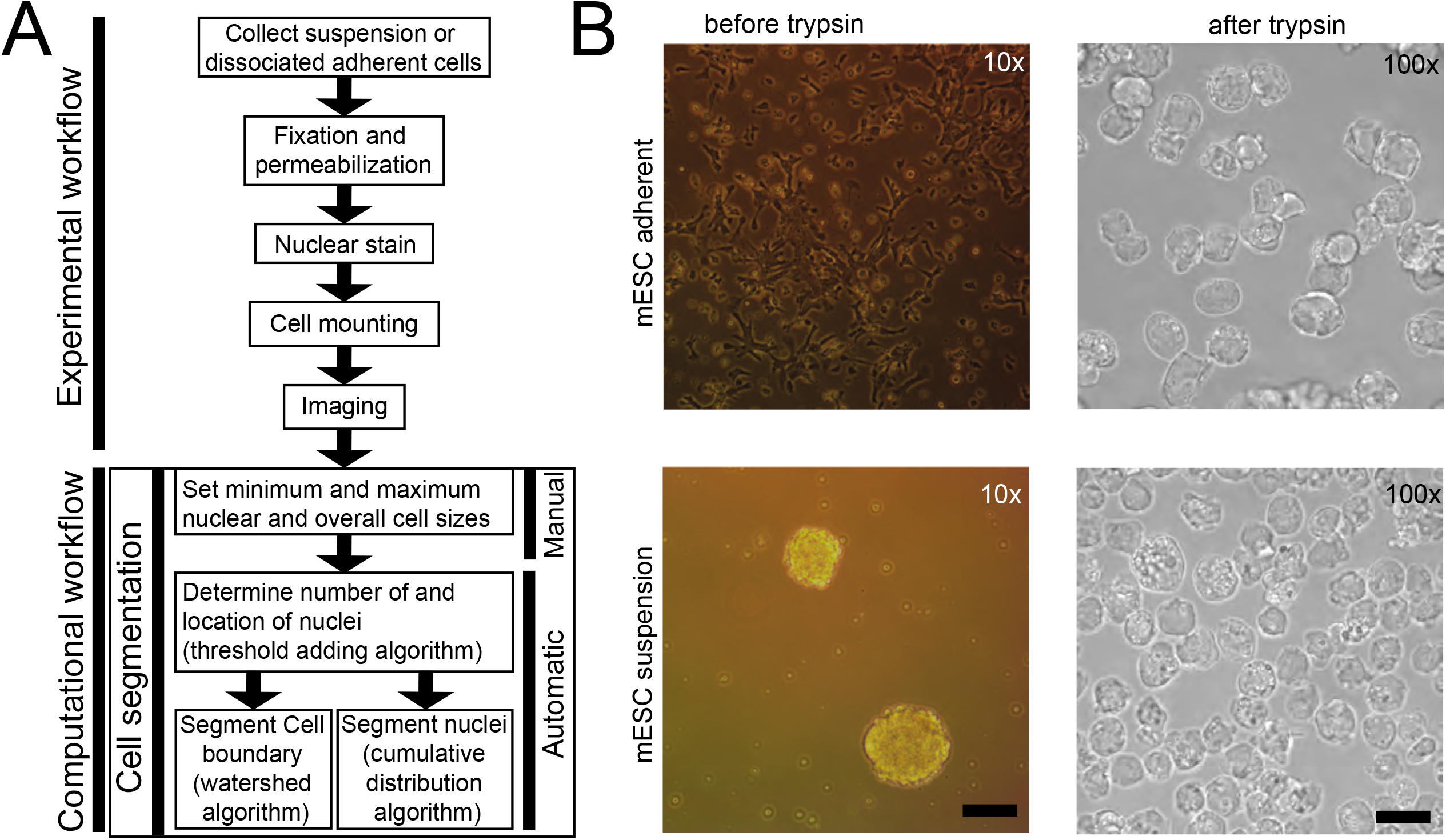
Sample preparation and computational requirements for automated cell segmentation. (**A**) Overview of experimental and computational workflow. Predefined minimum and maximum nuclear and cytoplasmic area for different cell types are selected and only need to be modified once for a specific cell type and imaging condition. (**B**) Cells can be adherent, in suspension, or exist as cell aggregates before trypsin treatment. After trypsinization and dissociation, single cells are imaged at high density on a microscope regardless if they were previously adherent or in suspension. Scalebar at 10x is 17.53 μm and at 100 × 5.07 μm.

## Results

Since we observed low-contrast cell boundaries in adherent cells or cell aggregates, while boundaries of single-cell suspensions tended to be more distinct, we hypothesized that suspension and single-cell dissociation could improve data quality for cells not already in single-cell suspensions. This idea became the first step in our CellDissect approach, in which adherent cells (Figure 1B, top) and cell aggregates (Figure 1B, bottom) are trypsinized to dissociate into single-cell suspensions. Cells are subsequently fixed with formaldehyde and permeabilized with ethanol, the DNA is stained with DAPI, and the cells are mounted on commercial microwell plates to ensure high cell density for imaging (Figure 1B, right). The CellDissect approach ensures that single fixed cells are in suspension and form round spheres that generate a strong refraction pattern from cell membranes upon widefield illumination (Figures 1B,3B). This characteristic is critical to identify the cellular boundary with CellDissect if cells are not naturally in a single-cell suspension, and it eliminates the need for a fluorescent cell boundary marker. Another advantage is that a standard widefield fluorescent microscope available in many cell biology facilities can be used without modifications. Our CellDissect approach is suitable if the scientific question does not require to know the exact cell shape in 3D or the cell-to-cell context in a cell culture plate or tissue. After image acquisition, the computational workflow in CellDissect (Figure 1A) consists of defining minimum and maximum nucleus and cell sizes for a specific cell type that can be determined by using a GUI. This step needs to be performed once for a specific cell type and microscope setup. After these parameters have been determined in a small number of cells, correct nuclear segmentation in 3D (Figure 2) is followed by cell boundary segmentation in 2D (Figure 3).

**Figure 2.**
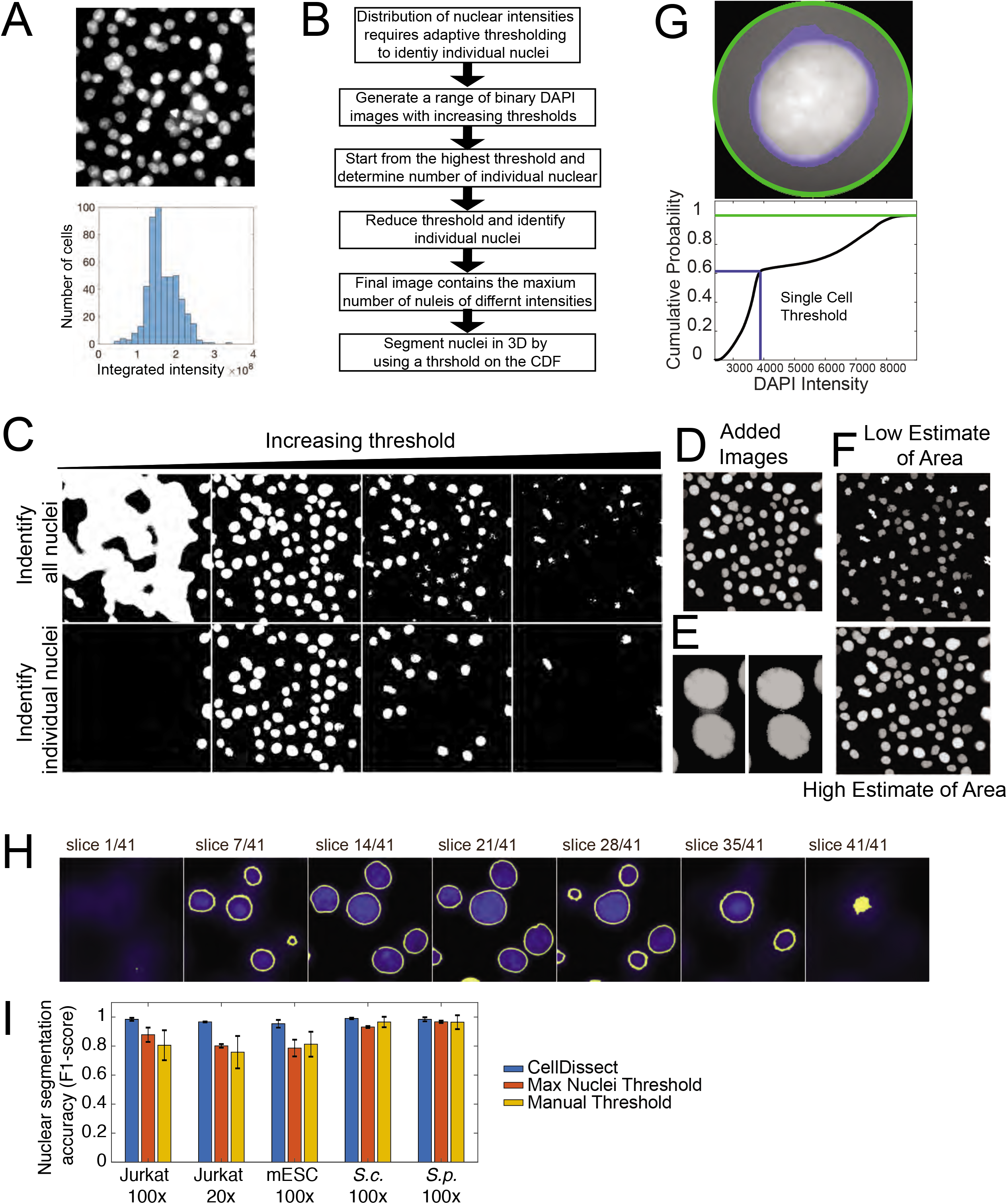
Automated nuclear segmentation of 3D image stacks. (**A**) Maximum projected cell nuclei with variable DAPI intensities (top). Distribution of integrated DAPI intensities within an image (bottom). (**B**) Workflow of the nuclear segmentation code after providing cell type-specific definitions of minimal and maximum nuclear area. (**C**) Increasing threshold result in an increase and then decrease in connected regions (top row). Binary images from the top are analyzed for connected regions within a cell type specific size range (bottom row). (**D**) Threshold binary images in the top row are added resulting in a pseudo image of large and small connected regions (Added Images). (**E**) Separation of connected objects before and after Low and High estimate of nuclear area. (**F**) Low and High estimate of nuclear area after watershed algorithm labels individual nuclei resulting in segmented nuclei. (**G**) Boundary of the nucleus is determined individually by radially integrating the fluorescent intensity from the outside (top, green circle). For each cell a threshold of 60% of the maximum cumulative fluorescent intensity (blue line) was chosen to robustly identify the nuclear boundary in 3D across all cell types and imaging conditions (bottom). (**H**) Nuclear segmentation in 3D from the bottom to top presented as series of images. (**I**) Nuclear segmentation accuracy (F1-score) quantification of 3D nuclear segmentation comparing fully automated adaptive threshold algorithm CellDissect (blue), maximum nuclei threshold algorithm (orange) and manual thresholding (yellow) in comparison to manual segmentation in four different cell lines and two different imaging magnifications. Mean and errors are computed from quantifying 3-4 images by three independent human experts.

**Figure 3.**
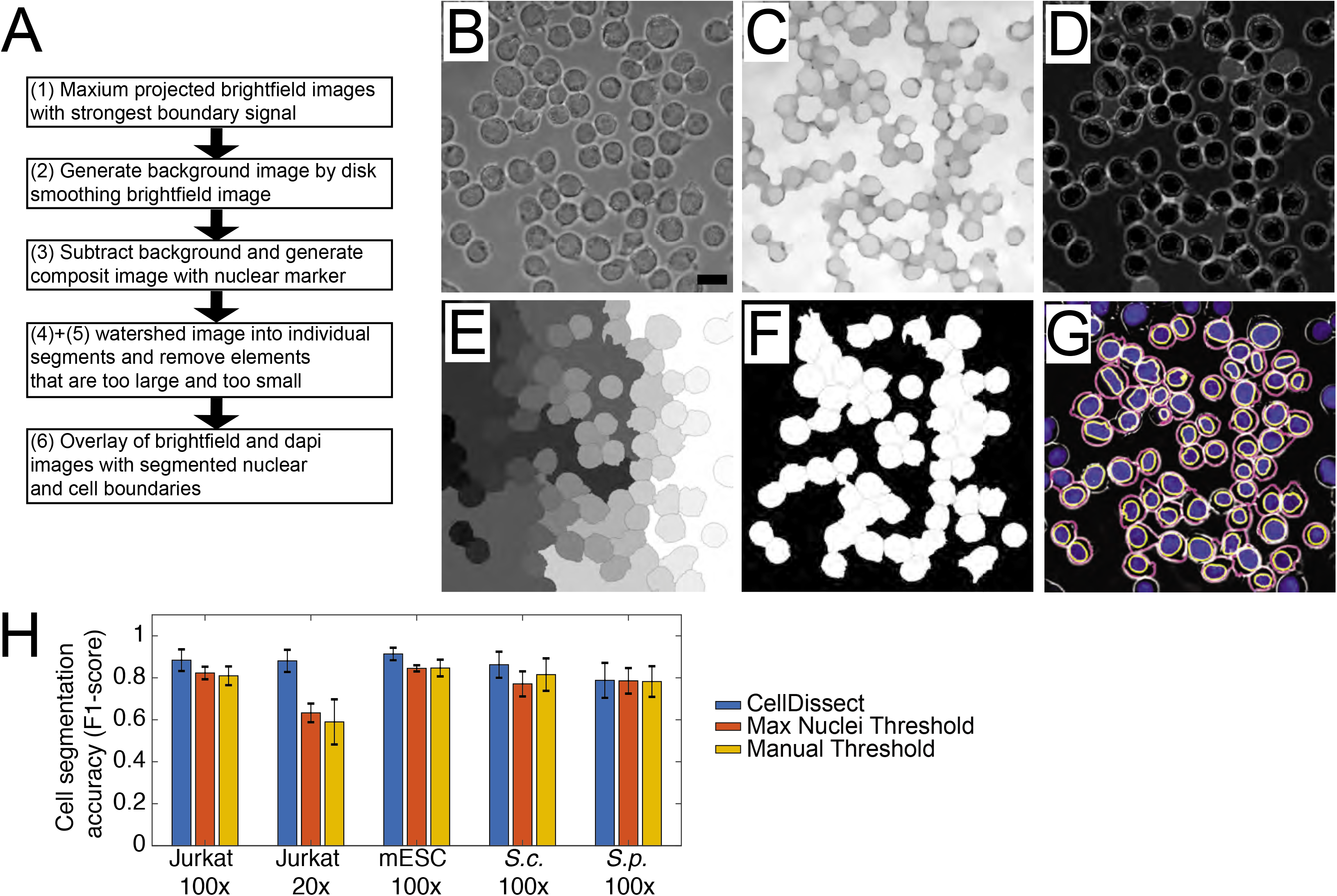
Automated cell boundary segmentation. (**A**) Workflow of the automated cell boundary segmentation. (**B**) Maximum projected widefield image using several images with clear cell boundaries is used to generate a **(C)** background image using disk smoothing. **(D)** After background subtraction, the maximum projected widefield image is overlaid with the segmented nuclei. **(E)** A watershed algorithm is applied resulting in segmented cells with cells on the image boundary removed **(F)**. **(G)** Overlay of widefield (grey), DAPI (blue), nuclear (yellow) and cytoplasmic (magenta) segmented mESC. Scalebar 13.3μm (**H**) Cell segmentation accuracy (F1-score) quantification of 2D cell segmentation comparing fully automated adaptive threshold algorithm CellDissect (blue), maximum nuclei threshold algorithm (orange) and manual thresholding (yellow) in comparison to manual segmentation in four different cell lines and two different imaging magnifications. Mean and errors are computed from quantifying the 3-4 images by three independent human experts.

### Nuclear segmentation in three dimensions

Traditionally, cells are segmented by using a fixed intensity threshold^16–19^. Although this has been sufficient to segment yeast cells, mammalian cells exhibit large variation in their nuclear DNA resulting in variable DAPI intensities (Figure 2A). If the goal is to precisely segment the nucleus, then the intensity of each individual nucleus needs to be considered. In order to ensure segmentation of cells with different DAPI intensities, in CellDissect we propose an adaptive thresholding approach (Figure 2B). Here we generate binary images from maximum intensity projected DAPI images with increasing thresholds and filter objects based on size (Figure 2C). This size filtering depends on the user-defined minimum and maximum nuclear size, which helps eliminate most instances of nuclei being blended together due to a low threshold (since it results in an abnormally large object) as well as noise or bright subsections of nuclei that are smaller than a complete nucleus (Figure 2D). To separate connected nuclei, we investigate each object to determine if removing layers of the added binary image results in separate nuclei fitting the size requirements (Figure 2E). This methodology allows one to identify the maximum number of individual nuclei in each image, and two different masks are generated during this process (Figure 2F). The first is a mask that results from removing the greatest number of layers while retaining the greatest number of nuclei, which becomes an underestimate of the nuclear area (Figure 2F, top). The second is a mask resulting from removing the least number of layers while still retaining the maximum number of nuclei, which results in a slight overestimate of the nuclear area for each nucleus (Figure 2F, bottom).

After the initial identification of the maximum number of nuclei, each nucleus is thresholded individually to determine the correct nuclear boundary (Figure 2G). The individual threshold is identified by utilizing the overestimate of nuclear area in the previous step to identify a threshold in the cumulative distribution of the DAPI signal for each nucleus independently. Although the slight overestimate of nuclear area often results in some noise being included in the initial nuclear determination, this is largely corrected by a subsequent processing step that removes all but the largest connected volume in 3D. Our CellDissect approach allows for automated and precise 3D nuclear segmentation (Figure 2H). Finally, we compare the accuracy of three nuclear thresholding algorithms that we developed and tested, which are (1) the adaptive thresholding algorithm CellDissect (blue bar), (2) an algorithm that uses a fixed threshold that maximizes the number of segmented nuclei (orange bar) and (3) a manually chosen fixed threshold (yellow bar), to the original images using human experts (Figure 2I). We scored false positive cells and nuclei as those that were detected computationally but not segmented correctly (>20% error). False negatives were cells or nuclei that were not identified computationally but were identified by a human expert. Applying the CellDissect approach to several cell types measured at different magnifications resulted in high accuracy (F1-score) in nuclear segmentation when compared to the ground truth of cell segmentation from several researchers (Figure 2I). The F1-score is defined as the harmonic average of precision and sensitivity. Precision shows how many of the computationally segmented objects are segmented correctly. Sensitivity shows how many of the total objects are segmented correctly. Fixed thresholds yielded good results in determining the number of *S. cerevisae (S.c.)* and *S. pombe (S.p.)* yeast nuclei per image. However, fixed thresholds performed extremely poor in identifying the correct number of nuclei in mammalian cells. We overcame this problem with our adaptive thresholding approach CellDissect (Figure 2C-E). In addition, defining an individual DAPI intensity threshold for each cell was essential to correctly segment the nuclear boundary in 3D (Figure 2G,H). In summary, we have demonstrated with CellDissect, for a range of different cell types and imaging modalities, that using multiple thresholds increases the number of nuclei identified and is essential to correctly identifying the number of mammalian nuclei (Figure 2I). In addition, individual thresholding of nuclei is essential to correctly identifying the nuclear boundary and ensuring 3D nuclear segmentation for all the different cell types.

### Cell boundary segmentation

To segment individual cells in 2D, our CellDissect pipeline consists of maximum projection of widefield images, background correction, composite image generation, watershed cell segmentation and object removal based on size resulting in segmented single cells (Figure 3A). First, we project several maximum contrast widefield images by maximum intensity to generate the cell outline as a white ring (Figure 3B). Next, we generate a background image from the cell widefield image through disk smoothing (Figure 3C). We then subtract this background image to enhance the contrast in the widefield image. Next, we generate a composite image of the processed widefield image and the nuclear mask (Figure 3D). This composite image is the input for a watershed algorithm (Figure 3E). Elements that are too big, too small or on the image boundary are removed (Figure 3F). The overlay of the processed widefield image, the DAPI image, and the segmented nuclear and cytoplasmic boundary is shown in Figure 3G.

To demonstrate the robustness and throughput of our approach, we applied our CellDissect pipeline to *S. cerevisae (S.c.)*, *S. pombe (S.p.)* yeast cells, mouse embryonic stem cells (mESCs) and human Jurkat cells with images taken at 100x magnification (Figure 4A). We also applied our CellDissect approach to mESCs and Jurkat cells imaged at 20x magnification. These results demonstrate qualitatively how our optimized experimental cell preparation protocol results in high quality and high cell density images at different cell magnifications, which is the basis for imaging large numbers of single cells. In addition, we quantified accuracy (F1-score) of our CellDissect approach in comparison to the ground truth that was independently generated from three human experts (Figure 3H). These results show quantitatively, that Cell Dissect outperforms the other approaches regardless of cell type and magnification. Finally, we outline how our CellDissect can be utilized on a single desktop computer or on a computing cluster that processes images in parallel to increase throughput (Figure 4B).

**Figure 4.**
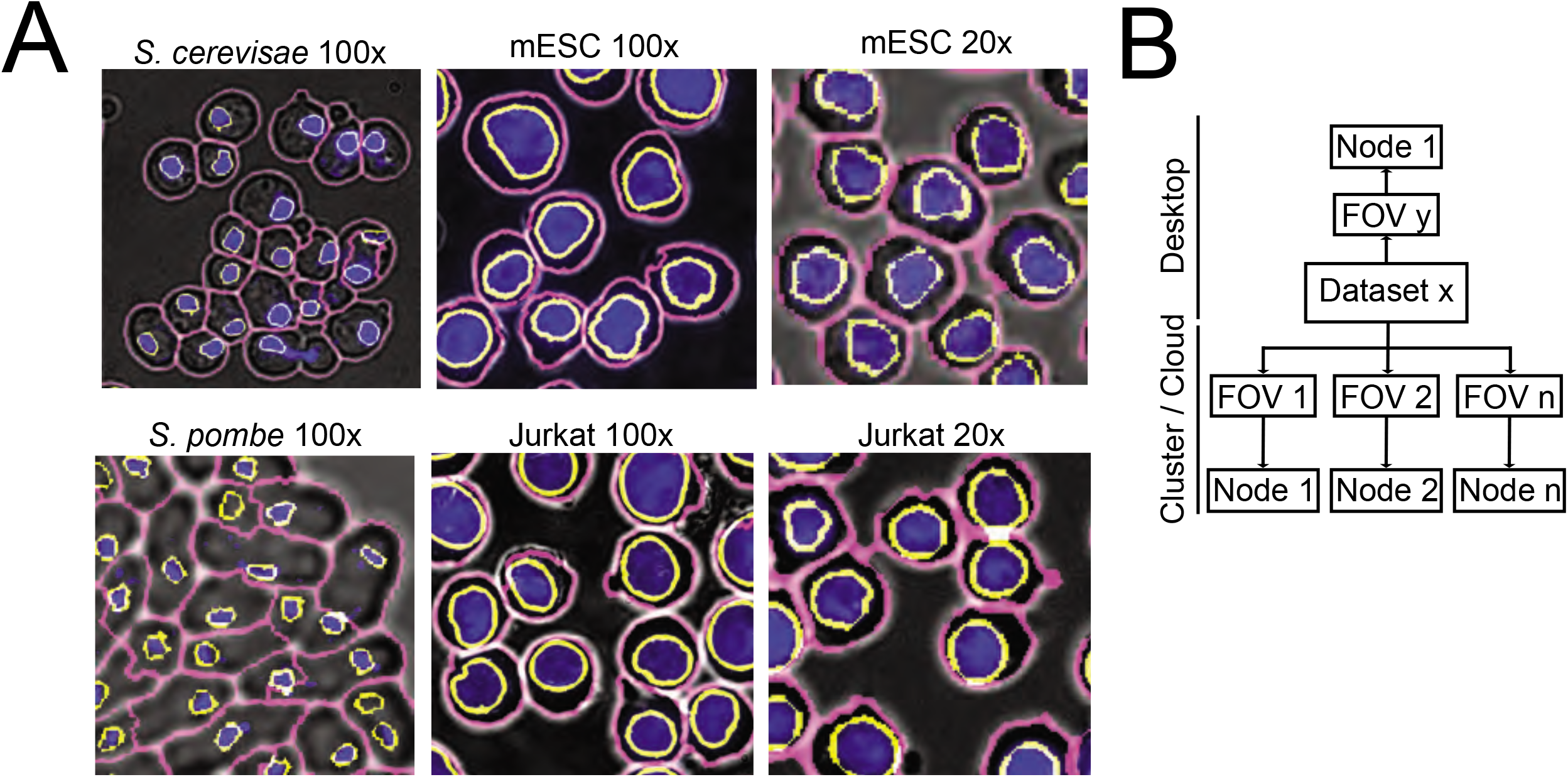
Application, implementation and performance of cell segmentation for different cell types from different organism imaged at low and high resolution. (**A**) Overlay of widefield (grey), DAPI (blue), nuclear (yellow) and cytoplasmic (magenta) segmented *S. cerevisiae*, *S. pombe*, mESC and Jurkat cells imaged at 100x and mESC and Jurkat cells imaged at 20x. (**B**) Image processing can be performed on a desktop computer or on a computing cluster for each individual image stack resulting in significant throughput.

## Discussion

We present CellDissect, a combined experimental and computational workflow to identify individual cells and segment their nuclei in 3D at high accuracy, at high cell density and with high throughput. In cases where cells are not already in a single-cell suspension and knowledge about cell morphology and nearest neighbor interaction is not required, dissociating cells into single-cell suspensions greatly improves data quality, uniformity and throughput (Figure 1)^22^. This is because cells form spheres in suspension, generating a strong diffraction pattern and allowing these cells to be imaged in widefield with high contrast cell boundaries (Figure 1B). Generating cell suspensions circumvents the problem of low contrast cell boundaries in adherent cells as well as the problems arising from three-dimensional cell aggregates. In addition, cell segmentation is independent of the density of imaged cells. With this optimized experimental workflow in CellDissect, that results in high contrast widefield images, we then improved nuclear segmentation by taking into consideration variable nuclear straining intensities due to differences in DNA content (Figure 2A, B). In CellDissect, we developed an adaptive thresholding algorithm that consists of two steps. In the first step, we apply a range of thresholds to identify all the nuclei in the image (Figure 2C-F). In the second step, we threshold each nucleus independently to account for the DAPI intensity differences, resulting in 3D segmented nuclei of high quality (Figure 2G, H). A quantitative assessment by human experts of nuclear segmentation confirmed CellDissect’s high accuracy, outperforming manually or automatically-chosen single threshold approaches (Figure 2I). CellDissect works very well for different types of cells imaged at different magnifications without the need for a large amount of training data or crowd source approaches^20^.

The nuclei that were initially determined are used as seeds to segment cell boundaries (Figure 3). Qualitative comparison between widefield images and the segmented images demonstrate the high accuracy of CellDissect’s image segmentation (Figure 3H). We then applied our CellDissect approach to other cell lines imaged at different magnifications (Figure 4A) and continued to observe a very high accuracy in nuclear and cell segmentation of all cell types using the F1-score metric in comparison to the ground truth generated by several humans (Figure 3H). CellDissect is written in a modular manner that is amenable to processing images either on a desktop computer or in parallel on a computing cluster to increase throughput (Figure 4B)^20^. The settings for both applications are generated through a GUI that can be adapted to different cell lines and microscope modalities^23–25^. These results show highly accurate nuclear and cell segmentation without the need for large training data sets. In addition, high accuracy in nuclear and cell segmentation is achieved by only using widefield imaging and DAPI stained nuclei. Recently developed cell segmentation approaches using deep learning indicate that cell segmentation is possible at high quality but most often requires very large data sets of images and significant hardware infrastructure to infer model parameters^3,9,14^.

In summary, by utilizing the experimental workflow in CellDissect, we prepare high quality singlecell suspensions that are subsequently imaged at high density, resulting in high quality images. These images are then analyzed by the computational workflow in CellDissect, that consists of reliable segmentation of nuclei and cells at high accuracy for a range of cell types and magnification without the need for large training data sets.

## Materials and Methods

### Experimental methods

#### Cell culture

The *Saccharomyces cerevisiae* (*S. cerevisiae*) strain *BY4741* (*MATa his3*Δ*1 leu2*Δ*0 met15*Δ*0 ura3*Δ*0*) was used and cultured as previously described^26^.

The *Saccharomyces pombe* (*S. pombe*) strain *972h*- was used. Three days before the experiment, *S. pombe* cells were streaked out on a YES (0.0002% each of adenine, histidine, leucine, lysine, uracil (w/v), 0.25% yeast extract) + 3% glucose plate from a glycerol stock stored at −80°C. The day before the experiment, a colony from the YES plate was inoculated in 5 ml YES + 3% glucose media (pre-culture) and grown at 32°C. After 6-12h, the optical density (OD) of the pre-culture was measured and the cells were diluted in new YES + 3% glucose media to reach an OD of 0.8 the next evening.

For imaging at 20x, the mouse embryonic stem cell (mESC) cell line 16.7 (Lee and Lu 1999) was grown with 1 million seeded cells on 75 cm^2^ tissue culture flasks with vented caps (Falcon 353110) gelatinized with EmbryoMax 0.1% Gelatin Solution (Millipore ES-006-B) for 30 minutes at 37C and plated with 2 million C57Bl/6 mouse embryonic fibroblasts as feeder cells (Gibco A34960) and with serum+LIF media composing of: DMEM with high glucose (Life Technologies 11960-044), 15% ES Cell qualified FBS (Gibco 16141-061), 25 mM HEPES (Gibco 15630-030), 1x MEM NEAA (Life technologies 11140-050), 1x (100 U/mL) Penicillin-Streptomycin (Gibco 15140-122), 100 μM 2-mercaptoethanol (Life Technologoies 21985-023), 500 U/mL LIF (EMD Millipore ESG1106), 1x GlutaMAX™ (Gibco 35050-061), and 1x (1 mM) sodium pyruvate (Gibco 11360-070). Cells were grown at 37°C in a 5% CO_2_ humidity-controlled environment for two passages before experiments.

For imaging at 100x, mESCs were thawed onto an MEF plate with conditioned media serum+LIF media as described for the 20x. The next day, media was changed with 2i media composed of: DMEM with high glucose (Life Technologies 11960-044), 25 mM HEPES (Gibco 15630-030), 0.5x MEM NEAA (Life technologies 11140-050), 1x (100 U/mL) Penicillin-Streptomycin (Gibco 15140122), 100 μM 2-mercaptoethanol (Life Technologies 21985-023), 1000 U/mL LIF (EMD Millipore ESG1106), .25x GlutaMAX™ (Gibco, Catalog#: 35050-061), and 1x (1 mM) sodium pyruvate (Gibco 11360-070), 20 μg/mL human insulin (Sigma I9278-5ML), 1 μM (Sigma PD0325901), 3 μM (Sigma CHIR99021), 1000 U/mL LIF (EMD Millipore ESG1107). After three days, the cells were passaged onto a plate gelatinized with 0.1% gelatin without feeders and grown for another passage.

Jurkat, Clone E6-1 (ATCC^®^ TIB-152™), cells were cultured at 0.5-1* 10^6 cells/ml in RPMI 1640 media (Corning, Catalog#: 15-040-CV) containing 10% Heat inactivated FBS (Gibco 16140-071), 1x Penincillin-Streptomycin (Gibco, Catalog#: 15140-122) and 1x GlutaMAX™ (Gibco 35050-061) at 37°C in a 5% CO_2_ humidity controlled environment.

#### Cell Fixation

*S. cerevisiae* were fixed in 4% formaldehyde as previously described^26^.

*S. pombe* cells were fixed with 1% formaldehyde for 15 minutes at room temperature, quenched with 150 mM glycine for 5 minutes at room temperature and set on ice for 5 minutes afterwards. They were then washed twice with 2x SSC and then permeabilized with 70% ethanol overnight.

mESCs were dissociated after washing with 1x PBS using accutase when cultured in 2i media and 0.05% trypsin when in serum + LIF media. The cell suspension was centrifuged for 5 minutes at 200 g, washed with 1x PBS, and then fixed for 8-10 minutes at room temperature with a 3.7% formaldehyde solution in 1x PBS. The cells were washed twice with 1x PBS and then permeabilized with 70% ethanol at 4°C for at least one hour.

Jurkat cells were fixed in their media described above with 2% formaldehyde for 10 minutes at room temperature. They were centrifuged for 3 minutes at 1000x*g* and then permeabilized with 100% methanol on ice.

#### DAPI staining

The washing and staining procedure was the same for all cells and has been previously described^21^, though their centrifugation times and speeds were different and matched what was described above.

#### Microscopy

Cells were imaged with epifluorescence as previously described^26^. Yeast cells were on 75 × 25 mm Corning microslides (2947-75×25) with 22×22 glass coverslips (12-542-B). Mammalian cells were on the ibidi 15 μ-Slide Angiogenesis (81506) in wells coated with 0.01% poly-D-lysine (Cultrex 3439-100-01) for 10 minutes.

### Computational methods

#### Size and slice determination (GUI)

The directory and details of images to be analyzed were input into the GUI. The first image from the directory was loaded, and ellipses were drawn to approximate the maximum and minimum nuclear and cellular sizes within the image. Individual slices for the cell boundary were looked at by eye, and a range was determined for when there were bright boundaries in the image.

#### Nuclear Determination

For adaptive nuclear thresholding, 100 evenly-spaced thresholds between the minimum intensity and maximum intensity were calculated. For each threshold from the 10^th^ to the 100^th^, a binary image of DAPI signal above the threshold was generated. MATLAB function “bwareaopen” filtered out objects that were too small or large based on the previously defined minimum and maximum nuclear sizes. All the binary images were added together, and each individual object was labeled using “bwlabeln”. It was determined for each individual object if more objects (within the size ranges) would result from removing layers from the binary images added together. Two new binary images resulted: 1) an image with a lower estimate of the nuclear area in the maximum intensity (dapi_label). For this, the maximum number of layers were removed that still had the maximum number of objects that fit the size restrictions. 2) An image with a higher estimate of the nuclear area in which the minimum number of layers were removed for maximum object number (dapi_label_low1).

For comparison, a single threshold was automatically determined. Thresholds were calculated, binary images were generated, and objects were filtered by size as described above. The threshold resulting in the maximum number of nuclei was applied, and this binary image was saved as both dapi_label and dapi_label_low1 for further processing steps.

As another comparison, a manually determined DAPI threshold was also used. A DAPI intensity threshold was picked by eye to identify the maximum number of individual nuclei (not blended together) in the image. This threshold was applied, and the resulting binary image was saved as both dapi_label and dapi_label_low1.

#### Nuclear Segmentation

Each object in dapi_label_low1 was investigated individually by determining a circle around the center of the object with a radius 1.3 times what would be expected from a circle matching the maximum nucleus size. The maximum intensity projection from the DAPI channel was determined inside this circle. Other nuclei in dapi_label_low1 within the circle other than the nucleus in the center were ignored. The cumulative distribution function (cdf) of DAPI intensity inside the circle was determined, and the area of the nuclear object in dapi_label_low1 was divided by the area of the circle and subtracted from 1 to find a threshold for the nucleus for the cdf. The corresponding intensity value was determined and applied to the circle in 3D (now a cylinder) to determine the nucleus. The other nuclear objects in dapi_label_low1 within the cylinder were again ignored. The objects in each slice were filled in with the MATLAB function “imfill”, and only the largest connected volume in 3D was kept. Nuclei too close to the image border were removed.

#### Cell segmentation

A maximum intensity projection of the slices defined by the user was used to start. A background image was generated with a disk smoothing filter over this image based on the minimum cell size. The background image was then subtracted from the maximum projected image. This image was then combined with processed DAPI image corresponding to the low estimate of nuclear area (dapi_label) using morphological reconstruction and cells were segmented with a watershed algorithm. After image segmentation, segmented elements that were too small or large elements were removed. Cells too close to the borders were also removed.

#### Quantification of Precision, Sensitivity, and Accuracy

To quantify sensitivity and precision, images from the DAPI and transmitted light channels as well as the segmented nuclei and cells were loaded and displayed in one overlaid image. Experts in our lab labeled each image for false positives and false negatives for both the nuclear and cellular segmentation. A false negative was defined as a lack of segmentation for an object. A false positive was defined as a segmented object with greater than 20 percent error in its area (positive or negative). For instances where two objects were determined as one object, one false positive and one false negative was counted. Objects on the edge of the image were not counted. True positives (TP) were calculated by subtracting the number of false positives from the number of segmented objects determined by the program. Sensitivity was calculated by dividing the number of true positives by the number of false negatives (FN) plus true positives: TP/(FN+TP). Precision was calculated by dividing the number of true positives by the number of false positives (FP) plus true positives: TP/(FP+TP). The accuracy was calculated as the F1-score which is the harmonic average of the precision and sensitivity: 2(Precision x Sensitivity)/(Precision + Sensitivity).

## Acknowledgements

BK, GL, AT, RV, and GN are supported by NIH DP2 GM11484901, NIH R01GM115892, and Vanderbilt Startup Funds. Additionally, BK was supported by T32GM008320, RV was supported by T32LM012412 and AT is supported by an AHA predoctoral fellowship (Award#: 18PRE34050016). Vanderbilt ACCRE computing cluster is supported by NIH S10OD023680. We would like to thank J.T. Lee for sharing mESC 16.7 cell line and K. Gould for sharing *S. pombe 972h* strain.

## Author contribution

BK and GN developed the image processing pipeline. BK, GL, RV and AT generated the images and analyzed the data. BK, AT, RV cultured the cells. BK, AT, RV and GN wrote the manuscript.

## Data availability

The datasets and computer codes generated and/or analyzed during the current study are available in the public dropbox repository: https://www.dropbox.com/sh/egb27tsgk6fpixf/AADaJ8DSjab_c0gU7N7ZF0Zba?dl=0

